# Putative role of PV and SOM interneuron subtypes in working memory

**DOI:** 10.1101/2021.05.31.446430

**Authors:** Pritish Patil, Mikhail Katkov, Ofer Yizhar, Misha Tsodyks

## Abstract

Working memory is an essential human trait required for all cognitive activities. Our previous model from Mongillo et al. (1), Mi et al. (2) uses synaptic facilitation to store traces of working memory. Thus memories can be maintained without persistent neural activity. A critical component of this model is a central inhibition which prevents multiple item representations from being active at the same time. We know from experimental studies that multiple genetically-defined interneuron subtypes (e.g. PV, SOM) with different excitability and connectivity properties mediate inhibition in the cortex. The role of these sub-types in working memory however is not known. Here we develop a modified model with these interneuron subtypes, and propose their functional roles in working memory. We make concrete testable predictions about the roles of these groups.

## Introduction

Working memory (hereafter WM) can be defined as “The ensemble of components of the mind that hold a limited amount of information temporarily in a heightened state of availability for use in ongoing information processing” (3). WM is an essential human cognitive process required for all cognitive activities.

The standard model of working memory postulates cells that show persistent firing during delay period encode working memory. Mongillo et al. (1) have suggested a plausible alternative mechanism for working memory. In this model, the trace of working memory is maintained by the facilitation of synapses. Synapses, once facilitated, maintain this state for time periods of upto a second without need for reactivation. Mi et al. (2) explain the limited capacity of WM in this model, which emerges out of temporal multiplexing of items. They show this model can simultaneously maintain several working memory items without interfering with one another. Strong recurrent inhibition in the model which prevents two items from being active simultaneously, thus preventing them from interfering with one another. Thus inhibition in this model plays an important role in the operation of working memory. However, inhibition in the cortex comes from multiple sources. In the cortex, inhibitory neurons can be roughly partitioned into distinct non-overlaping subtypes : PV, SOM, and VIP. (4, 5). These subtypes are known to have specific excitation and connectivity properties. They have been implicated to play distinct roles in different parts of the cortex. Experimental studies have demonstrated that these distinct interneuron subtypes play different roles in working memory (6–9); their particular roles however are not well understood.

Our aim here is to propose the possible role of interneurons in working memory. To this end, we try to create and simulate a minimal model that illustrates the different functions of interneuron types. Further, we make specific experimentally testable predictions of this model.

## Results

### Model description

Here we consider an extended version of the model from (1, 2) of working memory with memory items encoded by neuronal groups with short-term synaptic plasticity (STP).

Our model consists of multiple non-overlapping groups of excitatory neurons (Pyr) each encoding a single memory item. We assume that each group of neurons acts as a single unit which can be characterized by a single ‘firing rate.’ We incorporate two types of interneuron groups: PV and SOM, into our model. We have only one group of PV neurons and a SOM neuron group assigned to every Pyr group.

**Fig. 1** shows the connectivity diagram of our model. The model has two types of connections, facilitating and simple. In a simple connection, the postsynaptic current is a direct reflection of the presynaptic firing rate. In facilitating synapses, the postsynaptic current depends on history of presynaptic firing rates. We use the Tsodyks-Markram model (10) for facilitating connections. We use dynamic synapses only where modeling considerations suggest synapses with STP play a role. We also neglect some known synaptic connections to reduce the complexity of the model.

**Fig. 1.**
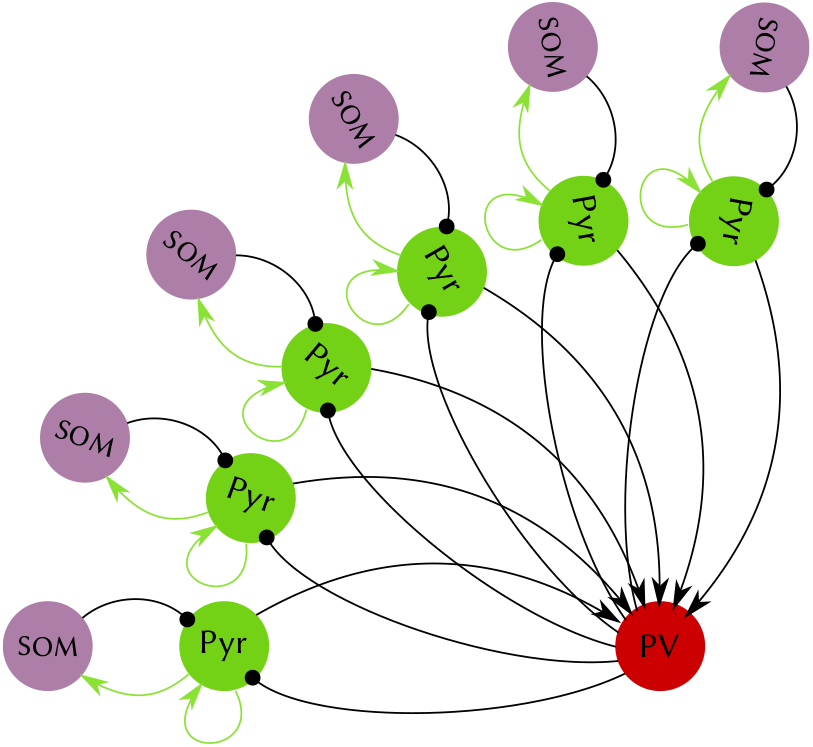
Schematic description of model. Our system consists of 3 types of neural populations. Pyramidal (Pyr), Parvalbumin expressing (PV), and Somatostatin expressing (SOM) neurons. Each population has excitatory (lines with arrows) or inhibitory (lines with dot at the end) synapses. The excitatory synapses can be simple (current proportional to firing rate, blacl lines) or dynamic (current depends on short-term synaptic plasticity, STP, green lines)

The Pyr groups have facilitating recurrent connections to themselves. They also have simple excitatory connection to the PV group. The PV group in turn has inhibitory connection to all the Pyr groups. Each Pyr group has a facilitatory excitatory connection to a single SOM group assigned to it, which reciprocally inhibits the Pyr group.

The dynamics of the model are described by the following equations:

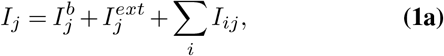

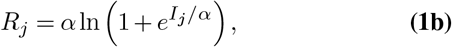

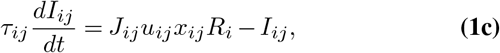

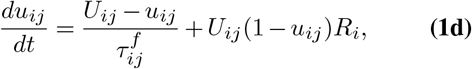

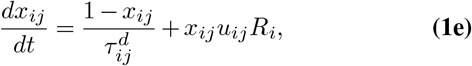

The total synaptic current *I_j_* for the group *j* is the sum of all the synaptic, external and baseline currents Eq. (1a). The firing rate *R_j_* of a group is a non-linear function of its total input current Eq. (1b). For each synapse from group *i* to group *j*, we have one equation for the corresponding synaptic current *I_ij_* Eq. (1c). If the synapse is dynamic, we have two additional variables : *u_ij_* and *x_ij_* for each synapse which represent “probability of release” and “amount of synaptic resources” respectively. The equations Eq. (1d) and Eq. (1e) describe the facilitation and depression dynamics respectively. In case of a simple synapse *u_ij_* and *x_ij_* variables are both fixed at 1. See Tsodyks et al. (10) for more details on short term synaptic plasticity model.

Experimental evidence suggests that PV interneurons provide strong and fast inhibition to Pyr cells which can implement the winner-take-all mechanism(11). Therefore we propose that the inhibition in the model in Mi et al. (2) is from PV interneurons. Further, the inhibition from SOM is relatively weak as compared to that from PV. (12). SOM interneurons innervate the dendrites of the post synaptic cells. They are known to receive facilitating excitatory inputs (13). Thus we propose that SOM interneurons are involved in locally controlling behavior of the working memory circuits.

## Simulations with PV and SOM

In order to understand the role of SOM interneurons in this system, we compare results of simulations with Mi et al. (2) model where the SOM interneurons are absent.

In the model without the SOM the memories once loaded stay loaded until the end of the simulation. An example of the dynamics is presented in **Fig. 2**a. In this simulation the first stimulus was loaded at 11 s to allow the network to reach steady state. Then we load 4 memories with a gap of 100 ms between them. The groups of neurons that represent each of these loaded items display a short interval of high firing rate, subsequently called population spike. These population spikes recur, without external stimulation, and thus constitute working memory.

**Fig. 2.**
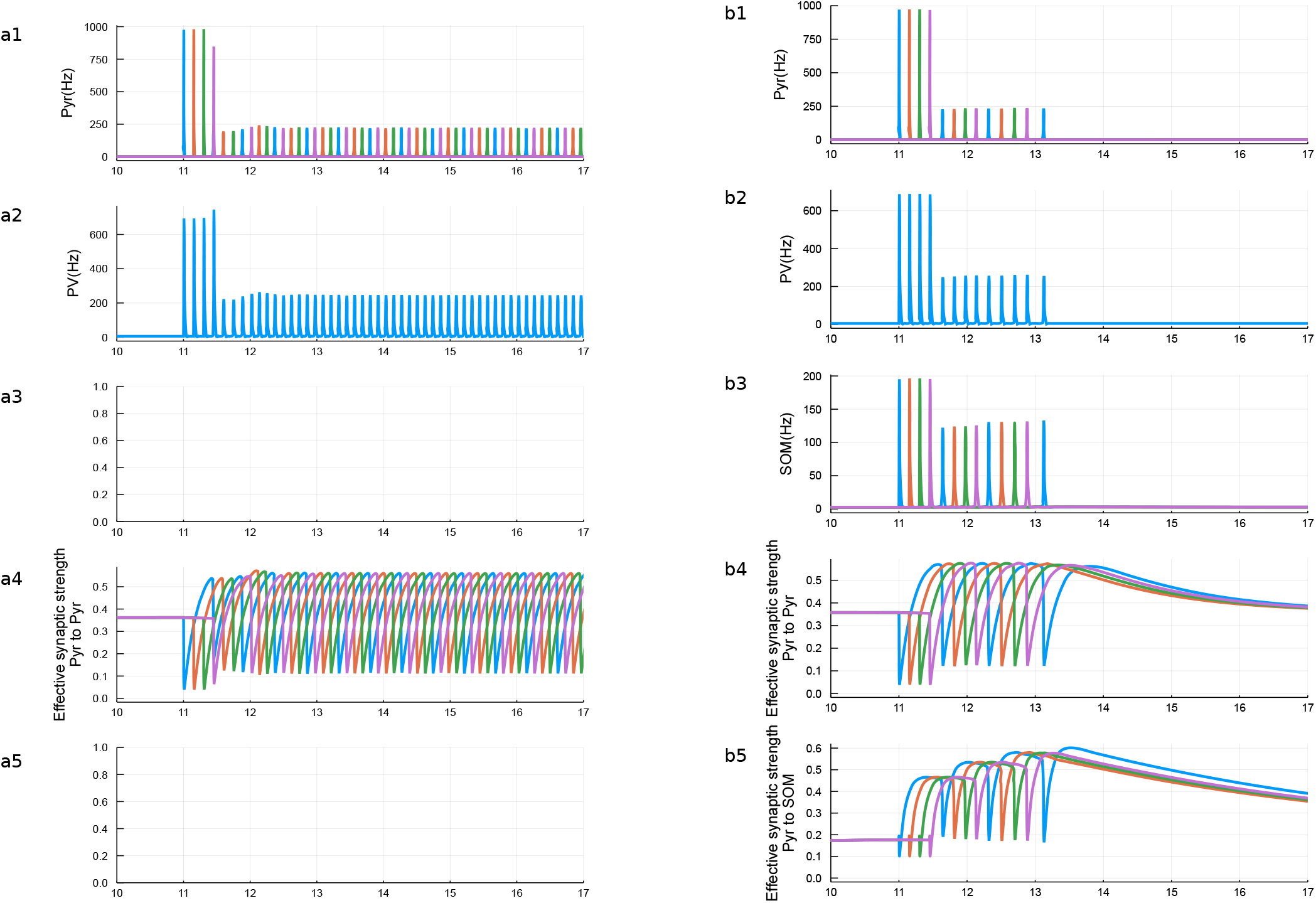
Comparison of the model with and without SOM. (a) Shows the model without SOM (as in Mi et al. (2)). (b) Model with SOM. Top row shows firing rate of Pyramidal neurons. One can observe that in the model without SOM neurons memories once loaded are oscillating for the rest of simulation, whereas in the presence of SOM population spikes disappear after few cycles. Second row shows activity of PV neurons. Third row shows activity of SOM neurons that does not exist in Mi et al. (2) model, and in our model show gradual increase of firing rate due to facilitatory connections from pyramidal neurons which are shown in bottom row.

In contrast, the system with SOM is capable of unloading (i.e. forgetting) loaded memories automatically. **Fig. 2**b shows an example of such dynamics. The mechanism of this forgetting is dependent on the facilitating synapses from Pyr to SOM. In the beginning, when these synapses are not facilitated, SOM neurons don’t play a big role and thus the system behaves similar to the model without SOM. Items are loaded into working memory and the corresponding group of neurons have population spikes periodically. These population spikes provide facilitating input to SOM. The net inhibition received by the Pyr groups increases as facilitation increases. Eventually the increased inhibition overcomes the excitation and the oscillations stop. **Fig. 2**b5 shows the effective synaptic strength of inhibition from SOM increases every cycle, which causes oscillations to stop after a few cycles (**Fig. 2**b1).

**Fig. 3** shows that the amount of time these oscillations persist can be controlled by the baseline input to SOM interneurons. This immediately suggests a mechanism for emergence of and local control of transience of working memory.

**Fig. 3.**
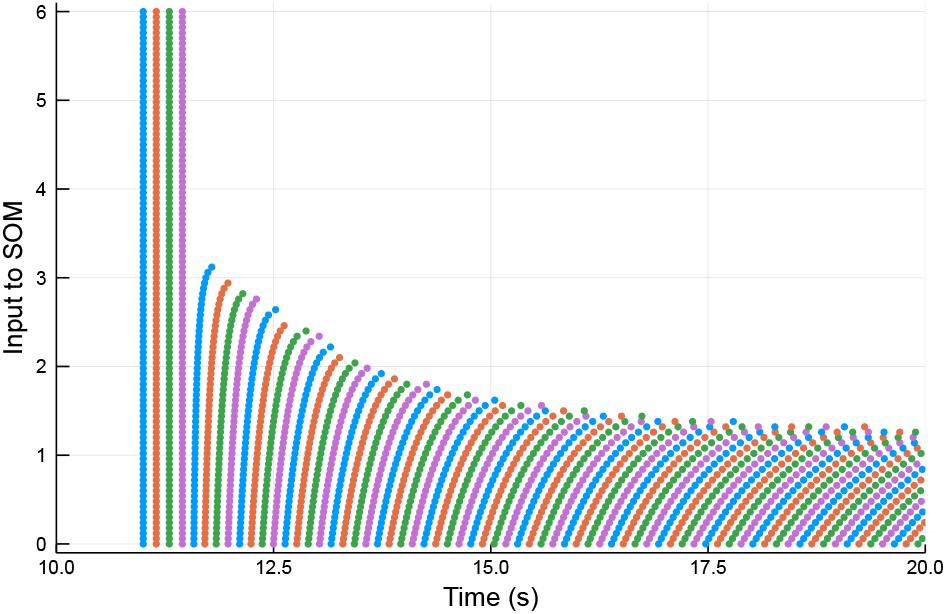
The effect of different baseline input currents to SOM group. Each row of dots corresponds to a simulation with a particular value of baseline input to SOM. Each dot is a population spike. Below a threshold input to SOM (~ 1.3) the memories are stable similar to those in Mi et al. (2). Above this threshold, in a range we get control over the duration these memories remain loaded. With a higher input to SOM cells (~ 3.0) the loaded memories fail to spontaneously reactivate at all.

## Discussion

In this study we designed a neural network model of working memory with the particular emphasis on the differential roles of PV and SOM interneurons. We suggest that the PV expressing interneurons are responsible for the “winner-takes-all” dynamics of working memory. A similar role has been suggested in other parts of the brain recently by other studies (14). Further, we show that SOM expressing interneurons and their facilitating excitatory inputs are responsible for the transient nature of working memory. (6) report the effect of optogenetic manipulation of SOM and PV interneurons during a spatial working memory task. They indicate that external optogenetic excitation of SOM group during the delay period affects performance, but only for long delay periods, and for very high intensity external excitation. This is consistent with our findings, as added baseline current to SOM shortens the length of the transient oscillations responsible for maintaining working memory. They also demonstrate that PV neurons were active throughout the delay period, a finding that is consistent with the idea that PV neurons are responsible for the “winner-take-all” dynamics.

Kim et al. (6) have also indicated that SOM neuron firing was informative about the target (i.e. item in memory), but the PV neuron firing was not. That the PV neurons firing was not selective to the targets is further consistent with the idea that PV neuronal group acts as a single unit, and all the neurons response to global activity and play a role in E-I balance and gain control.

We know from experimental studies that PV interneurons have strong synapses onto pyramidal neurons and have been found to mainly synapse onto soma or the axonal initial segment (4). Therefore they are thought to control the output of pyramidal neurons, consistent with the winner-takes-all function assigned to them in our model. PV interneurons have been implicated for this role in the Dentate Gyrus (14).

There is more known information about specific connectivity: PV neurons have strong self connections, and very strong autapses (15). They also have gap junctions with other PV neurons, and tend to act synchronously(11). Moreover, SOM interneurons are known to avoid making synapses onto other SOM interneurons (4). These and other features will be implemented in further versions of the model.

In summary, we have made concrete and experimentally testable predictions about the role of the different interneuron subtypes in working memory.

**Table 1.** Table listing parameters used in all the simulations:

|  | From | To | Parameter | Value |
| --- | --- | --- | --- | --- |
| Facilitating synapse | Pyr | Pyr | $J$ | 8.0 |
| | Pyr | Pyr | $U$ | 0.3 |
| | Pyr | Pyr | $\tau^d$ | 0.3 |
| | Pyr | Pyr | $\tau^f$ | 1.5 |
| | Pyr | Pyr | $\tau$ | $8 \times 10^{-3}$ |
| Facilitating synapse | Pyr | SOM | $J$ | 2.9 |
| | Pyr | SOM | $U$ | 0.1 |
| | Pyr | SOM | $\tau^d$ | 0.1 |
| | Pyr | SOM | $\tau^f$ | 5.0 |
| | Pyr | SOM | $\tau$ | $8e \times 10^{-3}$ |
| Simple synapse | Pyr | PV | $J$ | 1.75 |
| | Pyr | PV | $\tau$ | $8 \times 10^{-3}$ |
| Simple synapse | PV | Pyr | $J$ | -1.1 |
| | PV | Pyr | $\tau$ | $8 \times 10^{-3}$ |
| Simple synapse | SOM | Pyr | $J$ | -0.375 |
| | SOM | Pyr | $\tau$ | $15 \times 10^{-3}$ |
| Baseline Inputs | - | Pyr | $I^b$ | 3.0 |
| | - | PV | $I^b$ | -2.2 |
| | - | SOM | $I^b$ | 2.2 |
| Non linearity | - | - | $\alpha$ | 1.5 |

## ACKNOWLEDGEMENTS

This work was supported by Israeli Science Foundation grant 1657/19.

## Supplementary Note 1: Methods

Parameters used for simulations are in the Table 1:

Numerical simulations were performed in Julia language (16) using ODE solving package “DifferentialEquations.jl” (17), implementing of “DP8” Hairer’s 8/5/3 adaption of the Dormand-Prince 8 method Runge-Kutta method, a variable step size Runge-Kutta solver with 7th order interpolant. The relative and absolute errors were controlled to be less than 10^−6^.

## Bibliography

1. Gianluigi Mongillo, Omri Barak, and Misha Tsodyks. Synaptic theory of working memory. Science, 319(5869):1543–1546, mar 2008. ISSN 1095-9203. doi: 10.1126/science.1150769.

2. Yuanyuan Mi, Mikhail Katkov, and Misha Tsodyks. Synaptic correlates of working memory capacity. Neuron, 93(2):323–330, jan 2017. ISSN 08966273. doi: 10.1016/j.neuron.2016.12.004.

3. Nelson Cowan. The many faces of working memory and short-term storage. Psychonomic Bulletin & Review, 24(4):1158–1170, aug 2017. doi: 10.3758/s13423-016-1191-6.

4. Robin Tremblay, Soohyun Lee, and Bernardo Rudy. Gabaergic interneurons in the neocortex: from cellular properties to circuits. Neuron, 91(2):260–292, jul 2016. ISSN 08966273. doi: 10.1016/j.neuron.2016.06.033.

5. Henry Markram, Maria Toledo-Rodriguez, Yun Wang, Anirudh Gupta, Gilad Silberberg, and Caizhi Wu. Interneurons of the neocortical inhibitory system. Nature Reviews. Neuroscience, 5(10):793–807, oct 2004. ISSN 1471-003X. doi: 10.1038/nrn1519.

6. Dohoung Kim, Huijeong Jeong, Juhyeong Lee, Jeong-Wook Ghim, Eun Sil Her, Seung-Hee Lee, and Min Whan Jung. Distinct roles of parvalbumin- and somatostatin-expressing interneurons in working memory. Neuron, 92(4):902–915, nov 2016. doi: 10.1016/j.neuron.2016.09.023.

7. Andrew J Murray, Marta U Woloszynowska-Fraser, Laura Ansel-Bollepalli, Katy L H Cole, Angelica Foggetti, Barry Crouch, Gernot Riedel, and Peer Wulff. Parvalbumin-positive interneurons of the prefrontal cortex support working memory and cognitive flexibility. Scientific Reports, 5:16778, nov 2015. doi: 10.1038/srep16778.

8. Tsukasa Kamigaki and Yang Dan. Delay activity of specific prefrontal interneuron subtypes modulates memory-guided behavior. Nature Neuroscience, 20(6):854–863, jun 2017. doi: 10.1038/nn.4554.

9. Atheir I Abbas, Marina J M Sundiang, Britt Henoch, Mitchell P Morton, Scott S Bolkan, Alan J Park, Alexander Z Harris, Christoph Kellendonk, and Joshua A Gordon. Somatostatin interneurons facilitate hippocampal-prefrontal synchrony and prefrontal spatial encoding. Neuron, 100(4):926–939.e3, nov 2018. ISSN 08966273. doi: 10.1016/j.neuron.2018.09.029.

10. M Tsodyks, K Pawelzik, and H Markram. Neural networks with dynamic synapses. Neural Computation, 10(4):821–835, may 1998. doi: 10.1162/089976698300017502.

11. Hua Hu, Jian Gan, and Peter Jonas. Interneurons. fast-spiking, parvalbumin+ GABAergic interneurons: from cellular design to microcircuit function. Science, 345(6196):1255263, aug 2014. doi: 10.1126/science.1255263.

12. Sabine Krabbe, Enrica Paradiso, Simon d’Aquin, Yael Bitterman, Chun Xu, Keisuke Yonehara, Milica Markovic, Jan Gruendemann, Francesco Ferraguti, and Andreas Luthi. Adaptive disinhibitory gating by VIP interneurons permits associative learning. BioRxiv, oct 2018. doi: 10.1101/443614.

13. Aurélie Pala and Carl C H Petersen. In vivo measurement of cell-type-specific synaptic connectivity and synaptic transmission in layer 2/3 mouse barrel cortex. Neuron, 85(1): 68–75, jan 2015. doi: 10.1016/j.neuron.2014.11.025.

14. Claudia Espinoza, Segundo Jose Guzman, Xiaomin Zhang, and Peter Jonas. Parvalbumin+ interneurons obey unique connectivity rules and establish a powerful lateral-inhibition micro-circuit in dentate gyrus. Nature Communications, 9(1):4605, nov 2018. ISSN 2041-1723. doi: 10.1038/s41467-018-06899-3.

15. Charlotte Deleuze, Gary S Bhumbra, Antonio Pazienti, Joana Lourenço, Caroline Mailhes, Andrea Aguirre, Marco Beato, and Alberto Bacci. Strong preference for autaptic self-connectivity of neocortical PV interneurons facilitates their tuning to *γ*-oscillations. PLoS Biology, 17(9):e3000419, sep 2019. doi: 10.1371/journal.pbio.3000419.

16. Jeff Bezanson, Alan Edelman, Stefan Karpinski, and Viral B Shah. Julia: A fresh approach to numerical computing. SIAM review, 59(1):65–98, 2017.

17. Christopher Rackauckas and Qing Nie. Differentialequations. jl–a performant and feature-rich ecosystem for solving differential equations in julia. Journal of Open Research Software, 5(1), 2017.

